# Dopaminergic modulation of human inter-temporal choice: a diffusion model analysis using the D2-receptor-antagonist haloperidol

**DOI:** 10.1101/2020.02.13.942383

**Authors:** Ben Wagner, Mareike Clos, Tobias Sommer, Jan Peters

## Abstract

The neurotransmitter dopamine is implicated in diverse functions, including reward processing, reinforcement learning and cognitive control. The tendency to discount future rewards in value over time has long been discussed in the context of potential dopaminergic modulation. Here we examined the effect of a single dose of the D2 receptor antagonist Haloperidol (2mg) on temporal discounting. Our approach extends previous human pharmacological studies in two ways. First, we applied state-of-the-art computational modeling based on the drift diffusion model to comprehensively examine choice dynamics. Second, we examined dopaminergic modulation of reward magnitude effects on temporal discounting. Drift diffusion modeling revealed reduced temporal discounting and substantially faster non-decision times under Haloperidol. Temporal discounting was substantially increased for low vs. high reward magnitudes, but this magnitude effect was largely unaffected by Haloperidol. These results were corroborated by model-free analyses as well as modeling via more standard approaches using softmax action selection. We previously reported elevated caudate activation under Haloperidol in this sample of participants, supporting the idea that Haloperidol elevated dopamine neurotransmission, e.g. by blocking inhibitory feedback via presynaptic D2 autoreceptors. The present modeling results show that during inter-temporal choice, this leads to attenuated temporal discounting and increased response vigor (shorter non-decision times).

## Introduction

Future rewards are discounted in value^1^ such that humans and many animals prefer smaller-sooner (SS) rewards over lager-but-later (LL) rewards, a process referred to as temporal or delay discounting. Steep temporal discounting of reward value is associated with a range of maladaptive behaviors ranging from substance-use-disorders^2^, attention-deficit hyperactivity disorder^3^ and obesity^4^ to behavioral addictions such as gambling disorder^5^. Because alterations in temporal discounting are associated with such a variety of disorders, temporal discounting has been suggested to constitute a transdiagnostic process^6,7^ with relevance for many psychiatric conditions.

The neurotransmitter dopamine (DA) plays a central role in the reinforcing effects of drugs of abuse as well as in mechanisms underlying core processes of addiction^8^. The rodent literature has relatively consistently shown increased temporal discounting following a reduction in DA neurotransmission and conversely a decrease in temporal discounting following moderate increases in DA^9^. However, the corresponding human literature is small and considerably more heterogeneous^9^. For example, de Wit et al.^10^ found that acute administration of d-amphetamine decreased impulsivity across a range of tasks including temporal discounting, such that temporal discounting was reduced under d-amphetamine. However, a later study did not replicate this effect^11^. Administration of the D2/D3 receptor agonist pramipexole did not affect measures of impulsivity in another study (n=10) from the same group^12^. In contrast, Pine et al.^13^ observed increases in temporal discounting following administration of the catecholamine precursor L-DOPA compared to placebo in healthy control participants (n=13), while the D2-receptor antagonist Haloperidol showed no significant effect on the discount rate. In a recent within-subjects study using L-DOPA in a substantially larger sample (n=87), L-DOPA, in contrast, had no overall effect on temporal discounting^14^. Rather, the effect of L-DOPA depended on baseline impulsivity, which the authors interpreted in the context of the inverted-U-model of DA effects on cognitive control functions^15^. Two recent studies have reported a reduction in discounting following administration of the selective D2/D3-receptor antagonist amisulpride^16^ as well as the D2 receptor antagonist metoclopramide^17^.

A similar heterogeneity is evident when considering tasks with close conceptual links to temporal discounting. For example, reinforcement learning (RL) is thought to depend on at least two systems, a model-based system that evaluates actions based on a model of the environment, and a model-free system that relies on the reinforcement of rewarded actions^18^. A conceptual relationship to temporal discounting is clear when considering that model-based planning is thought to rely on anticipatory mechanisms that compute future outcomes based on an environmental state-space, whereas such explicit forward planning is not required for model-free RL. While one study found no association between model-based RL and temporal discounting^19^, another large-scale online study observed the predicted positive association: participants with a stronger contribution of model-based planning to RL also showed reduced temporal discounting^20^. However, in contrast to temporal discounting, where (as mentioned above) elevation of DA levels via the catecholamine precursor L-DOPA lead to more impulsive decision-making and steeper discounting^13^ (though results in a larger sample were considerably more mixed^14^), L-DOPA instead *increased* reliance on model-based RL in controls^21^ and Parkinson’s disease patients^22^. However, a recent study in a substantially larger sample (n=65) did not replicate this overall effect of L-DOPA on model-based control^23^. Rather, in this study the effect of L-DOPA was restricted to a subset of participants with high working memory capacity.

Several task-related and contextual factors affect the degree of temporal discounting^1,24^. One well-replicated behavioral effect, the magnitude effect, refers to the observation that the rate of temporal discounting decreases with increasing reward magnitude^25^. In humans, this effect depends on lateral prefrontal cortex processing^26^. In rodents, effects of *d*-amphetamine on temporal discounting have been shown to depend on reinforcer magnitude, such that more pronounced effects of d-amphetamine were observed for large magnitude conditions^27^. However, a comprehensive analysis of dopaminergic effects on magnitude-related effects in human temporal discounting is still lacking.

Taken together, the literature on DA contributions to human temporal discounting using pharmacological approaches appears heterogeneous. Many previous studies^10,14,16^ have focused on temporal discounting tasks with low trial numbers that precluded comprehensive modeling of the behavioral data (but see^13^) and/or a detailed analysis of the magnitude-effect or response time (RT) distributions. In the present study, we therefore examined cognitive processes underlying temporal discounting and the magnitude-effect more comprehensively using a between-subjects double-blind placebo-controlled pharmacological approach using a low dosage (2mg) of the D2-receptor antagonist haloperidol. In separate memory tasks performed by our participants during fMRI, we have previously reported substantial increases in task-related dorsal striatal activation in the Haloperidol vs. Placebo group during trial onset both for recognition^28^ and associative recall^29^. This observation is compatible with a predominantly presynaptic effect of Haloperidol in the present study, leading to an overall increase in striatal dopaminergic signaling. Importantly, we extend previous pharmacological studies by applying a modeling framework for temporal discounting data that is based on a combination of standard discounting models with the drift diffusion model (DDM)^30–33^, allowing us to comprehensively examine drug effects on RT components related to both valuation and non-valuation related processes.

## Methods

### Participants

Fifty-four healthy participants were initially enrolled in the study. Twenty-seven participants were randomly assigned to each group [Placebo/Haloperidol]. Two participants from the Haloperidol group did not complete the temporal discounting task. Technical problems lead to working memory data loss from four participants (three from the Haloperidol, one from the Placebo group), but these participants were still included in the temporal discounting data analysis.

Following filtering of RTs (see below, the fastest and slowest 2.5% of trials were excluded per participant), we examined the individual RT histograms for each subject. This revealed that even after filtering, the three participants with the fastest minimum RTs (two from the Haloperidol group and one from the placebo group) still showed implausibly fast responses on a number of trials (minimum RTs of 2ms, 2ms and 234ms, see subjects 24, 25 and 41 in Supplemental Figure S1) such that the minimum RTs were substantially faster than those in the remaining participants (all min(RT) z-scores of −2.04, −2.04 and −1.7, see Supplemental Figure S2). These subjects were therefore excluded from further modeling.

We verified that there were no significant differences in demographic background in terms of age or baseline working memory capacity (see Table 1). All participants were screened by a physician for current diseases, current intake of prescription drugs or drugs of abuse. Only healthy subjects were allowed to participate. Potential side effects of the medication were monitored via multiple blood pressure and pulse measurements and evaluated via mood questionnaires. These analyses did not reveal significant group differences in terms of reported mood, side effects or physiological parameters, as reported in our previous study^29^. Before enrollment, participants provided informed written consent and all study procedures were approved by the local institutional review board (Hamburg Board of Physicians).

**Table 1.** Demographic and working memory data (mean +/− SD).

|  | Placebo | Haloperidol | Group comparison |
| --- | --- | --- | --- |
| Age | 24.4±3.4 | 23.4±2.5 | T<sub>47</sub>=1.20, p=.24 |
| Sex (m/f) | 7/18 | 6/18 | X<sup>2</sup><sub>1</sub>=.001, p=1 |
| WM-Baseline (Z-Score) | -.0453±.665 | .0943±.556 | T<sub>44</sub>= -.80, p=.43 |

### General procedure

The study consisted of two testing sessions performed on separate days. On the first day (T0) participants completed a background screening and a set working memory tasks (*see below*). On the second day (T1) participants received either placebo or haloperidol (2mg). In line with the pharmacokinetics of haloperidol^34^ testing on T1 was performed 5 hours after drug administration to ensure appropriate plasma levels of haloperidol. During the first 2.5hrs, participants where under constant observation and pulse as well as blood pressure levels were checked 30 minutes and 2 hours after drug administration. During the waiting period, participants filled out questionnaires on current mood and medication effects. Participants then completed a number of unrelated tasks during a functional magnetic resonance imaging (fMRI) scanning session (total scan-time 2.5 hrs.). Following scanning, they first completed the temporal discounting task outlined below, followed by a set of working memory tasks (digit span forward & backward, block span forward & backward, complex working memory span, see^29^ for detailed results).

### Temporal discounting task

Participants performed 210 trials of a temporal discounting task where on each trial they made a choice between a smaller-but-sooner (SS) reward available immediately, and a larger-but-later (LL) reward. SS and LL rewards were randomly displayed on the left and right sides of the screen, and participants were free to make their choice at any time. For half the trials, the SS reward consisted of 20€ and for the remaining trials the SS reward was fixed at 100€. These trials were presented randomly intermixed. LL options were computed via all combinations of a set of LL reward amounts (constructed by multiplying the SS reward with [1.01, 1.02, 1.05, 1.10, 1.20, 1.50, 1.80, 2.50, 2, 3, 4, 5, 7, 10, 13]) and LL delays (1, 2, 3, 5, 8, 30, 60 days), yielding 105 trials in total per magnitude condition. As is typically the case for temporal discounting tasks investigating magnitude effects^25^, all choices were hypothetical.

### Computational modeling

#### Temporal discounting model

We applied a simple single-parameter hyperbolic discounting model to describe how value changes as a function of delay^35,36^:

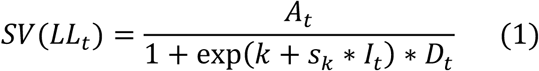

Here, *A*_*t*_ is the numerical reward amount of the LL option on trial *t*, *D*_*t*_ is the LL delay in days on trial *t* and *I* is an indicator variable that takes on a value of 1 for trials from the large-magnitude condition (SS amount = 100€) data and 0 for trials from the small-magnitude condition (SS amount = 20€). The model has two free parameters: *k* is the hyperbolic discounting rate from the large magnitude condition (modeled in log-space) and *s*_*k*_ is a weighting parameter that models the degree of change in discounting for small vs. large SS rewards (i.e., higher values in *s*_*k*_ reflect a greater magnitude effect^25^).

### Softmax action selection

Softmax action selection models the choice probabilities as a sigmoid function of value differences^37^:

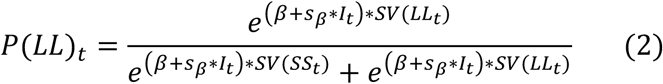

Here, *SV* is the subjective value of the risky reward according to Eq. 1 and *β* is an inverse temperature parameter, modeling choice stochasticity (for *β* = 0, choices are random and as *β* increases, choices become more dependent on the option values). *SV(SS*_*t*_) was fixed at 100 for the large magnitude condition and fixed at 20 for the small magnitude condition. I is again the dummy-coded condition regressor, and *s*_*β*_ models the magnitude effect on *β*.

### Temporal discounting drift diffusion models

To more comprehensively examine dopaminergic effects on choice dynamics, we additionally replaced softmax action selection with a series of drift diffusion model based choice rules. In the DDM, choices arise from a noisy evidence accumulation process that terminates as soon as the accumulated evidence exceeds one of two response boundaries. In the present setting, the upper boundary was defined as selection of the LL options, whereas the lower boundary was defined as selection of the SS reward.

RTs for choices of the SS option were multiplied by −1 prior to model fitting. We furthermore used a percentile-based cut-off, such that for each participant the fastest and slowest 2.5 percent of trials were excluded from the analysis. We then first examined a null model (DDM_0_) without any value modulation. Here, the RT on each trial *t* is distributed according to the Wiener First Passage Time (*wfpt*):

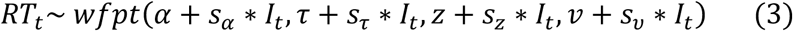

The parameter α models the boundary separation (i.e. the amount of evidence required before committing to a decision), τ models the non-decision time (i.e., components of the RT related to motor preparation and stimulus processing), *z* models the starting point of the evidence accumulation process (i.e., a bias towards one of the response boundaries, with *z*>.5 reflecting a bias towards the LL boundary, and *z*<.5 reflecting a bias towards the SS boundary) and *ν* models the rate of evidence accumulation. Note that for each parameter *x*, we also include a parameter *s*_*x*_ that models the change in that parameter from the high magnitude (SS=100) to the low magnitude (SS=20) condition (coded via a the dummy-coded condition regressor *I*_*t*_). As in previous work ^30,32,38^, we then set up temporal discounting diffusion models by making trial-wise drift rates proportional to the difference in subjective values between options. First, we set up a linear modeling scheme (DDM_lin_)^32^:

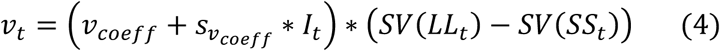

That is, the drift rate on trial *t* is calculated as the scaled value difference between the SS and LL rewards. As noted above, RTs for SS options were multiplied by −1 prior to model estimation, such that this formulation predicts SS choices whenever SV(SS)>SV(LL) (the trial-wise drift rate is negative), and predicts longest RTs for trials with the highest decision-conflict (i.e., in the case of SV(SS)= SV(LL) the trial-wise drift rate is zero). We next examined a DDM with non-linear trial-wise drift rate scaling (DDM_S_) that has recently been reported to account for the value-dependency of RTs better than the DDM_lin_^30,38^. In this model, the scaled value difference from Eq. 4 is additionally passed through a sigmoid function with asymptote *v*_*max*_:

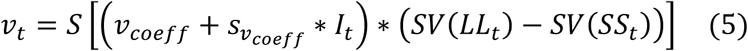

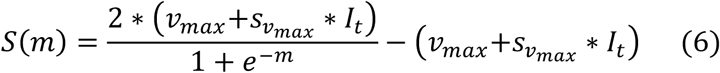

All parameters including *v*_*coeff*_ and *v*_*max*_ were again allowed to vary according to the reward magnitude condition, such that we included *s*_*x*_ parameters for each parameter *x* that were multiplied with the dummy-coded condition predictor *I*_*t*_ (see above).

### Hierarchical Bayesian models

Models were fit to all trials from all participants using a hierarchical Bayesian modeling approach with separate group-level distributions for all parameters for the Placebo and Haloperidol groups. Model fitting was performed using Markov Chain Monte Carlo as implemented in the JAGS software package^39^ (Version 4.3) using the Wiener module for JAGS that implements likelihood functions for the Wiener First Passage Time^40^ (see Eq. 3) in combination with R (Version 3.4) and the R2Jags package. For group-level means, we used uniform priors defined over numerically plausible parameter ranges. For all *s*_*x*_ parameters modeling condition effects on model parameters, we used Gaussian priors with means of 0 and standard deviations of 2. For group-level precisions, we used Gamma distributed priors (.001,. 001). We ran 2 chains with a burn-in period of 900k samples and thinning of two. 10k additional samples were then retained for further analysis. Chain convergence was assessed via the Gelman-Rubinstein convergence diagnostic *Ȓ* and values of 1 ≤ *Ȓ* ≤ 1.01 were considered as acceptable for all group-level and individual-subject parameters. Relative model comparison was performed via the Deviance Information Criterion (DIC), where lower values reflect a superior fit of the model^41^.

## Results

### Subjective and physiological drug effects

As reported in detail in our previous papers^28,29^, there were no significant group differences with respect to reported side-effects, subjective mood, heart rate or blood pressure relative to baseline. Likewise, groups did not differ with respect to the actual and guessed drug condition (Haloperidol vs. Placebo)^29^.

### Model free analysis of temporal discounting

Figure 1 depicts the overall response time (RT) distributions per group with choices of the LL option coded as positive RTs and choices of the SS option coded as negative RTs. RT histograms for each individual participant are shown in Supplemental Figure S1. As a model-free measure of temporal discounting, we examined proportions of LL choices as a function of group (placebo vs. haloperidol) and condition (100€ vs. 20€ reference reward). Raw proportions of larger-but-later choices are plotted in Figure 2. ANOVA on arcsine-square-root transformed proportion values with the within-subject factor *magnitude* (100€ vs. 20€ SS reward) and the between-subject factor *drug* (Placebo vs. Haloperidol) confirmed a significant magnitude effect (F_(1,47)_=96.86, p<.001) such that participants overall made more LL selections in the high magnitude condition. Furthermore, effects of drug (F_(1,47)_=3.47, p=.068) and drug x magnitude condition (F_(1,47)_=3.31, p=.075) showed trend-level significance.

**Figure 1.**
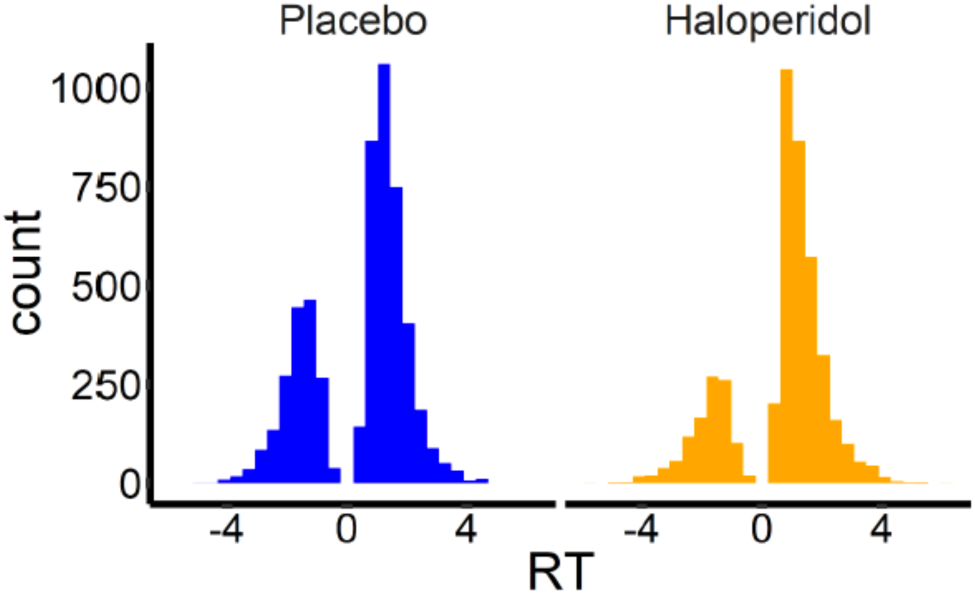
Overall response time (RT) distributions for the Placebo group (n=25) and the Haloperidol group (n=24). Negative RTs reflect choices of the smaller-but-sooner option, whereas positive RTs reflect choices of the larger-but-later option. It can be seen that participants in the Placebo group made numerically more SS selections than participants in the Haloperidol group.

**Figure 2.**
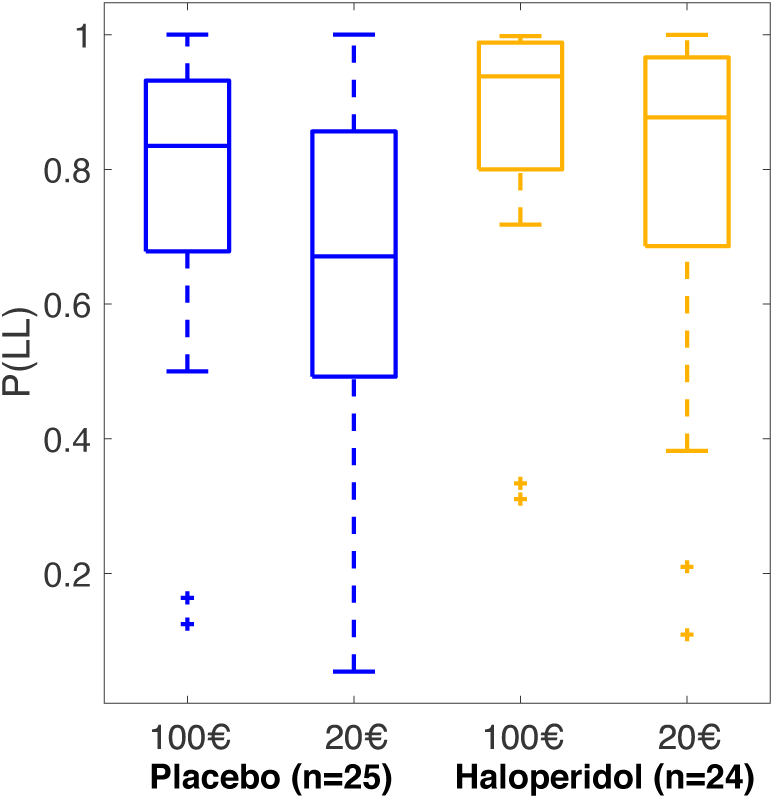
Proportion of larger-but-later choices per group and magnitude condition.

### Softmax choice rule

Results for a hierarchical Bayesian model with softmax choice rule are shown in Supplemental Figure S3. This analysis revealed a substantial magnitude effect on log(k) as well as an overall drug effect, such that log(k) was substantially lower in the Haloperidol group compared to the Placebo group.

### Model comparison

We next compared three versions of the drift diffusion model that varied in the way that they accounted for the influence of value differences on trial-wise drift rates based on the deviance information criterion (DIC)^41^. In each model, we included separate group level distributions for the two drug conditions (Haloperidol vs. Placebo). Furthermore, for each parameter *x*, we included a shift parameter *s*_*x*_ modeling the change in parameter *x* from the high magnitude condition (smaller-sooner reward =100€) to the low magnitude condition (smaller-sooner reward = 20€) (see methods section). These *s*_*x*_ parameters where modeled with Gaussian priors with means of zero (see methods section).

The null model (DDM_0_) assumes fixed drift rates independent of value. We compared this model to two variants of the DDM that assume that trial-wise drift rates depend on the value-differences between options, either in a linear fashion (DDM_lin_)^32^ or in a non-linear (sigmoid) fashion^30,38^. The data were best accounted for by the DDM with non-linear value-scaling of trial-wise drift rates (Table 2).

**Table 2.** Model comparison of three variants of the drift diffusion temporal discounting model. The data were best accounted for by a model including a non-linear mapping from trial-wise value-differences to drift rates (DDM_S_).

| Model | DIC | Rank |
| --- | --- | --- |
| DDM<sub>0</sub> | 20938 | 3 |
| DDM<sub>lin</sub> | 20817 | 2 |
| DDM<sub>s</sub> | 16744 | 1 |

### Overall group differences

We next examined overall group differences in model parameters for the baseline (smaller-sooner reward =100€) condition. Results are plotted in Figure 3 and Bayes Factors for all group comparisons are listed in Table 3. In both groups, there was a positive association between trial-wise drift rates and value differences, as the 95% HDI for the drift rate coefficient parameter *v*_*coeff*_ did not include 0 in either group (Figure 3b). Likewise, there was a slight bias towards the smaller-sooner option in both groups, as the 95% HDI for bias was <0.5 in both cases (Figure 3e).

**Table 3.** Summary of group differences in model parameters for the temporal discounting drift diffusion model. For each parameter, we report mean posterior group differences (M_diff_) and Bayes Factors (BF) testing for directional effects on both the baseline parameter in the 100€ condition (left columns) and on the magnitude effect on each parameter (right columns). Bayes Factors <.33 indicate evidence for placebo<haloperidol, whereas Bayes Factors >3 indicate evidence for placebo>haloperidol.

| Model parameter | Baseline |  | Magnitude effect |  |
| --- | --- | --- | --- | --- |
| | $M_{diff}$ | $BF$ | $M_{diff}$ | $BF$ |
| **Log(k)** | 2.26 | 77.9 | -.093 | .47 |
| **Drift rate coefficient ($v_{coeff}$)** | -.365 | .061 | .020 | 2.73 |
| **Non-decision time ($\tau$)** | .180 | 98.4 | -.0001 | .95 |
| **Boundary separation ($\alpha$)** | -.047 | .60 | .017 | 1.47 |
| **Starting point bias ($z$)** | -.004 | .74 | -.017 | .26 |
| **Drift rate maximum ($v_{max}$)** | .18 | 8.27 | .16 | 16.88 |

**Figure 3.**
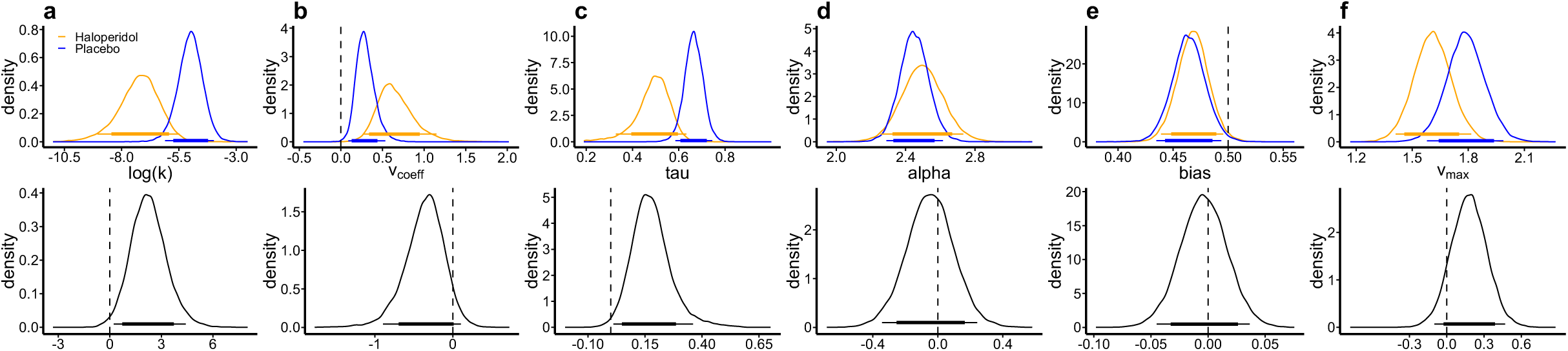
Posterior distributions per parameter (top row) and group differences (bottom row, placebo – haloperidol) for the baseline condition (smaller-sooner reward =100€). a: discount rate *log(k)*, b: drift rate coefficient *v*_*coeff*_, c: non-decision time *tau*, d: boundary separation *alpha*, e: starting point *bias*, f: maximum drift rate *v*_*max*_. Thin (thick) horizontal lines denote 95% (85%) highest posterior density intervals.

We furthermore observed substantially lower group-level discount rates log(k) in the haloperidol group compared to placebo, such that the 95% HDI of the posterior group difference in log(k) was >0 (Figure 3a). Interestingly, the non-decision time was likewise substantially lower in the Haloperidol group (Figure 3c), amounting to on average 180ms faster non-decision times.

### Magnitude effects on model parameters

We next turned to the effects of the magnitude manipulation on diffusion model parameters, that is, the change in each parameter in the 20€ condition compared to the 100€ baseline condition. Results are plotted in Figure 4 and Bayes Factors for all group comparisons are listed in Table 3. There was a substantial magnitude effect on log(k), such that discounting was steeper in the 20€ condition (Figure 4a). Interestingly, this pattern of results was not mirrored by in the magnitude effect on the starting point / bias parameter. Rather, the bias was shifted slightly in the direction of a neutral bias (0.5) in the low magnitude condition (Figure 4e) in both groups. An additional interesting observation is that the non-decision time was increased in the 20€ condition by on average around 30ms (Figure 4c).

**Figure 4.**
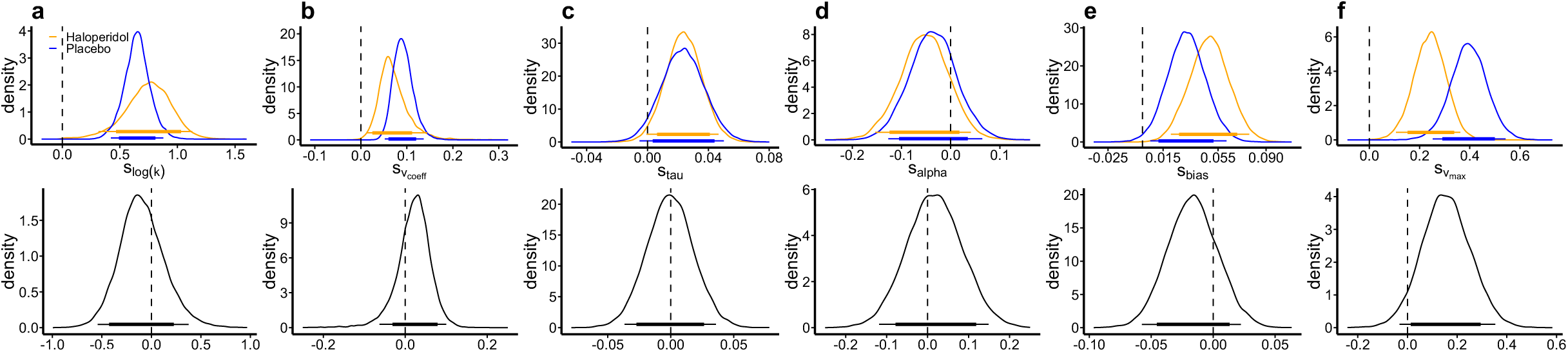
Posterior distributions of the change in each parameter in the 20€ condition compared to the 100€ baseline condition (top row) and corresponding group differences (bottom row, placebo – haloperidol) (a: discount rate *log(k)*, b: drift rate coefficient *v*_*coeff*_, c: non-decision time *tau*, d: boundary separation *alpha*, e: starting point *bias*, f: maximum drift rate *v*_*max*_). Thin (thick) horizontal line denote 95% (85%) highest posterior density intervals.

Both drift rate components (*v*_*coeff*_ and *v*_*max*_) were increased in the 20€ condition (Figure 4b and 4f). This overall effect can be attributed to the fact that in our model, these two parameters effectively scale the trial-wise value differences to the appropriate scale of the DDM^32^. Because average value differences spanned a smaller absolute range in the 20€ condition, this is compensated in the model by increasing the drift rate scaling parameters. There was, however, some evidence for a reduced magnitude effect on *v*_*max*_ (Figure 4f) in the Haloperidol group.

Difference distributions in the remaining model parameters were centered at zero, indicating no systematic group differences (Figure 4).

## Discussion

We investigated the effects of a single dose of the D2-receptor antagonist Haloperidol (2mg) on temporal discounting in a between-subjects study in a double-blind placebo-controlled setting. Comprehensive model-based analysis of choices and response times (RTs) revealed substantially smaller log(k) parameters in the Haloperidol group compared to Placebo, indicating reduced temporal discounting. We additionally observed a substantial reduction in non-decision times in the Haloperidol group.

We applied a recent class of computational models combining standard reinforcement learning and/or valuation models with the drift diffusion model^30,32,33,38^. Such a comprehensive analysis of RTs was not possible in most previous studies due to the specifics of task timing^13^ or the available trial numbers^14^ that precluded RT-based modeling^14,16^. The model comparison confirmed previous results^30,38^ such that the data were better accounted for by a model assuming a non-linear trial-wise scaling of the drift rate, as compared to a linear model or a null model without value modulation^30^. Importantly, we also fit our data using a more standard approach based on softmax action selection^37^. This reproduced the drug effects observed for the DDM-based modeling. Likewise, the general pattern of results was reproduced when examining balanced LL choice proportions as a model-free measure of temporal discounting. Both observations strengthen our confidence in the validity of the results observed for the DDM-based modeling approach. We have previously reported ^detailed parameter recovery analyses of the best-fitting DDMS model, which confirmed that^ both subject-level and group-level parameters recovered well^30^. Likewise, we previously reported detailed simulations confirming that key effects of value differences on RTs as well as decision consistency could be reproduced using that model^30^. Finally, the magnitude effect on temporal discounting is a well-replicated behavioral effect in the literature on temporal discounting ^25,26,42^. This effect was reproduced via the DDM-based modeling scheme in both groups, supporting previous findings^30^ that the effects of within-participant experimental manipulations of temporal discounting can be quantified and reproduced using diffusion modeling based approaches.

As highlighted in a recent review^9^, the human literature on dopamine (DA) contributions to impulsivity is characterized by considerable heterogeneity. The interpretation of these inconsistent findings in DA pharmacology are complicated by several additional factors. First, effects of dopaminergic drugs might depend on baseline DA availability^15^, such that the same drug might impair or enhance performance in different participants, according to an inverted-U-shaped function (or a different function that depends on the process that is examined^43^). Second, and more important to the present study, the action of D2-receptor antagonists are often interpreted in terms of a *reduction* in DA neurotransmission^13,44^. But such drugs might in fact *enhance* DA release by predominantly binding at presynaptic DA auto-receptors, at least at lower dosages^45^. Indeed, this has been observed in both animal studies ^46,47^ and human work^48^. Such an interpretation of D2-receptor antagonist effects in terms of a presynaptically-mediated elevation of DA release could potentially reconcile a number of conflicting results from human and animal studies. D2/D3 receptor antagonists have previously been shown to decrease temporal discounting in humans^16,17^ (with one study reporting a null result^13^), an effect that has in the animal literature been consistently linked to moderate *increases* in DA^9^.

Our findings help to resolve some of these conflicting results in several ways. First, our finding of reduced temporal discounting under Haloperidol is in line with two recent studies that reported reduced temporal discounting following administration of D2/D3 receptor antagonists ^16,17^. On the other hand, a reduction of temporal discounting following administration of Haloperidol was not observed in an earlier within-subjects study with n=13 participants^13^ that used a slightly lower dosage of 1.5mg (we used 2mg). As mentioned above, lower dosages of D2/D3 receptor antagonists might increase (rather than decrease) DA signaling^45^, an effect mediated by inhibitory feedback through presynaptic D2 autoreceptors^49^. This feedback is thought to lead to relatively specific enhancement of phasic (as opposed to tonic) DA signaling under Haloperidol^45^, a point that we return to below. However, we do acknowledge that such an interpretation is not general consensus in the cognitive literature on DA drug effects^13,44^.

Our results advance over previous findings regarding the role of D2/D3 receptor antagonists in temporal discounting^13,16,17^ in several ways. First, participants in the present study performed an unrelated memory task during fMRI directly prior to completing the temporal discounting task reported here. Those data revealed an overall main effect of drug condition on trial onset-related activity in caudate nucleus during recognition^28^ and associative recall^29^ (both effects were correlated^29^), such that participants from the Haloperidol group showed a substantially elevated dorsal striatal response in relation to stimulus onset, as compared to the placebo group. This neural read-out was obtained prior to the temporal discounting task, but both the fMRI and temporal discounting time points were well within the time of maximum Haloperidol plasma levels^34^. Although fMRI was measured during performance of different tasks, this observation is arguably more compatible with the idea that the dosage of Haloperidol applied in the present study increased (rather than decreased) striatal DA signaling. Similar neural evidence was lacking in most previous human pharmacological studies on DA contributions to temporal discounting ^10,12,16,17^. Second, the DDM-based modeling approach adopted in the present study (see above) allowed us examine the dynamics underlying decision-making much more comprehensively than previous human pharmacological studies^10,12–14,16,17^. In addition to the drug-effect on the discount rate log(k), diffusion modeling revealed a substantially shorter non-decision time in the haloperidol group that amounted to ≈180ms on average. Such a robust enhancement of lower-level motor and/or perceptual RT components is also more compatible with an increase rather than a decrease in DA transmission^50^ and resonates with previous findings regarding a dopaminergic enhancement of RT-based response vigor^51,52^. In support of this interpretation, augmentation of DA levels in Parkinson’s disease patients reduces temporal discounting^53^ and improves model-based reinforcement learning^22^. Finally, this interpretation of available human D2 receptor antagonist effects would also reconciliate the human and animal literature on acute dopaminergic effects on impulsivity^9^. Taken together, these considerations lead us to suggest that Haloperidol increased (rather than decreased) striatal DA neurotransmission, resulting in enhanced cognitive control (reduced discounting) and a substantial facilitation of motor responding (shorter non-decision times).

The question then arises by which mechanism Haloperidol might attenuate the impact of temporal delays on reward valuation. According to models of basal ganglia contributions to action selection^54^, the probability for selecting a given candidate action depends on the relative difference in activation between the direct (*go*) and the indirect (*nogo*) pathways. A similar striatal gating mechanism has been suggested to underlie the selection of representations for working memory maintenance and/or other higher cognitive functions supported by the prefrontal cortex^55^. By increasing phasic DA responses, Haloperidol might thus increase the signal-to-noise ratio in striatal value representations, thereby increasing the likelihood that objectively smaller and/or more delayed LL rewards gain access to processing in the prefrontal cortex. In Supplemental Figure S1, we illustrate how a change in the precision of LL value predictions can directly impact the overall likelihood of LL choices. Naturally, other modes of action are likewise conceivable. Frontal and striatal regions are interconnected via a series of loops that follow a dorsal-to-ventral organization^56^, and Haloperidol might impact functional interactions within these circuits^55^, e.g. related to top-down control of value representations^57–60^. A final possibility is that Haloperidol might have augmented cognitive control functions by directly augmenting processing in specific prefrontal cortex regions^59^. However, due to the much greater expression of D2 receptors in striatum compared to prefrontal cortex^61^, it is generally assumed that prefrontal action of D2 antagonists require substantially higher dosages than those applied in the studies examined here ^45,61^.

In a related line of work, individual differences in DA neurotransmission have been examined via positron emission tomography (PET) measures. Here, reductions in striatal D2 receptor density have repeatedly been associated with substance-use disorders^62,63^ (but not gambling disorder^64^), with the degree of decision-making impairments in clinical groups^65^ and with degrees of impulsivity in healthy control participants^66^ (but see^65^ for lack of evidence of an association between D2 receptor density and temporal discounting in healthy controls). The idea that reduced D2 receptor signaling is associated with both addiction and impulsivity is also supported by a large body of animal work^62^. But reconciliation of the human PET literature and the available pharmacological literature is again complicated by the fact that in particular low dosages of D2 antagonists such as Haloperidol might produce predominantly presynaptic effects, thereby increasing DA release, rather than producing a net reduction in overall D2 receptor signaling^45,47^.

The present study has a number of limitations that need to be acknowledged. First, we did not run a within-subjects design, which would have allowed us to account for individual-participant baseline parameters in the analysis of the drug effects. Second, this also precluded us from comprehensively analyzing potential modulatory influences of e.g. individual differences in working memory on the drug effects, which might play a role in the context of dopaminergic effects on temporal discounting^14^ and cognitive control more generally^15^. Finally, in the present study, all rewards were hypothetical. It is common practice in many studies on temporal discounting to e.g. randomly select a single trial following the testing session and pay out the selected option for real^67,68^. Given the high magnitude condition included in our design, this was not feasible. However, choice preferences for real and hypothetical outcomes in inter-temporal choice tasks show a very good correspondence^69^ and rely on similar neural circuits^70^.

Taken together, our data show that the D2 receptor antagonist Haloperidol attenuated temporal discounting and substantially shortened the non-decision time, as revealed by comprehensive computational modeling of choices and RTs using hierarchical Bayesian parameter estimation. In combination with Haloperidol increasing dorsal striatal responses in an unrelated task in our participants^29^, these data are best accounted for by a model in which low dosages of Haloperidol lead to an enhancement of phasic DA responses due to reduced feedback inhibition from D2 autoreceptors, leading to an augmentation of both low-level (non-decision time) and higher-level (temporal discounting) components of the value-based decision process.

## Acknowledgements

This work was supported by Deutsche Forschungsgemeinschaft (SO952/3-1 to T.S. and PE1627/5-1 to J.P.).

## Author contributions

MC, TS and JP designed the study. MC acquired the data. BW, MC and JP analyzed the data. BW performed the modeling. JP contributed analytical tools. BW and JP co-wrote the paper. MC and TS provided revisions.

## Supplemental Material

**Supplemental Figure S1.**
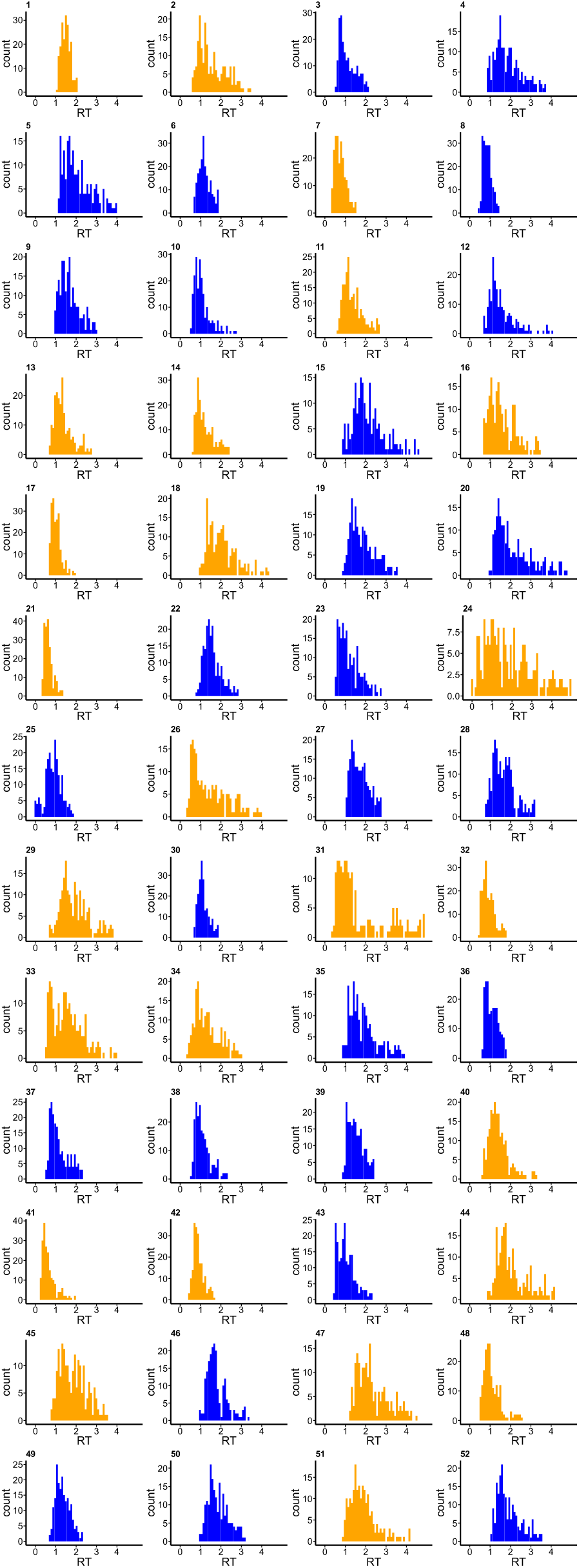
Response time (RT) histograms for each individual participant (placebo group: blue, haloperidol group: orange) following exclusion of each participants’ 2.5% fastest and 2.5% slowest trials. Subjects 24, 25 and 41 still had a number of implausibly fast trials even after filtering (see Supplemental Figure S2 and methods section) and were therefore excluded from modeling.

**Supplemental Figure S2.**
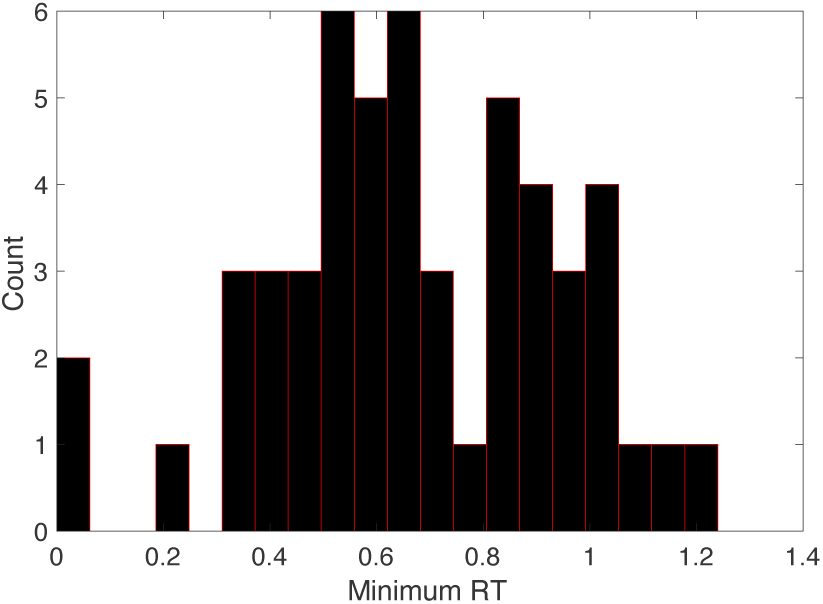
Histogram of minimum RTs across subjects following percentile-based trial filtering (i.e., following exclusion of each participants 2.5% fastest and 2.5% slowest trials). Still three participants (leftmost bars in the plot) had implausibly fast minimum RTs (z-scores < −1.65), corresponding to subjects 24, 25 and 41 from Supplemental Figure S1. These participants were excluded from modeling.

**Supplemental Figure S3.**
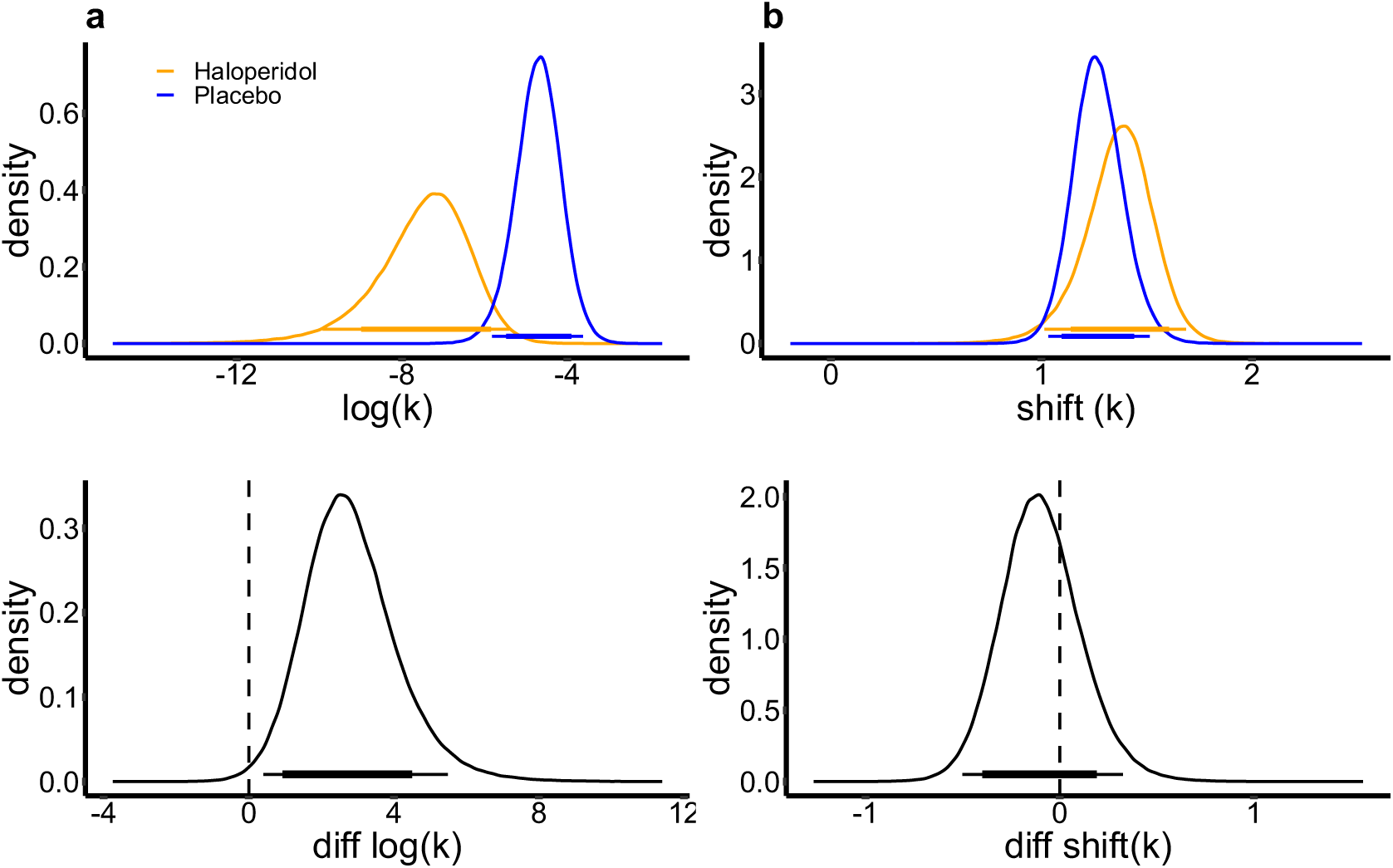
Modeling results from a hierarchical Bayesian Model with softmax choice rule (HAL - Haloperidol, PL - Placebo). Log(baseline_k) is the log(discount rate) from the high magnitude condition (smaller-sooner reward = 100€). Shift(k) is the change in log(k) from the high magnitude condition to the low magnitude condition (smaller-sooner reward = 20€). The thin (thick) horizontal lines denote 95% (85%) Highest Posterior Density Intervals.

**Supplemental Figure S4.**
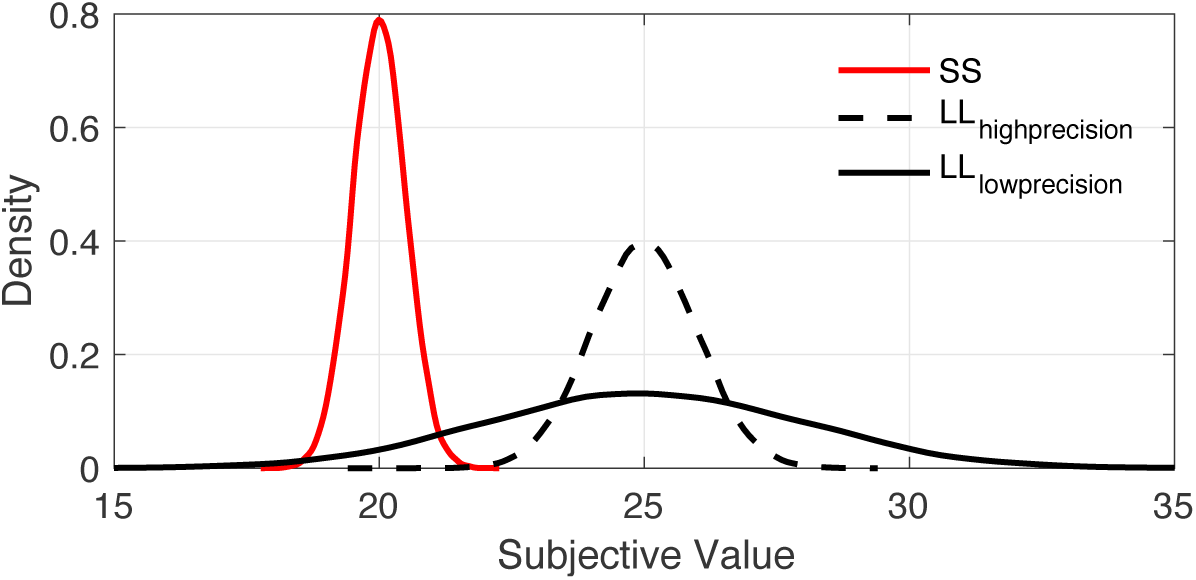
Illustration of how a sharpening of value representations can lead to a reduction in SS choices without shifting the mean of the LL value prediction. The value of a constant SS reward can be estimated with high precision (red distribution). In contrast, LL value predictions are assumed to be less precise than SS value predictions (black lines, the variance of these distributions is overall lower than for the SS reward). An LL value prediction with lower precision (solid black line) leads to a greater likelihood of SS choices even if the value prediction has the same mean, as illustrated by the larger overlap between red and solid vs. dashed black distributions.

